# Profiling proteins involved in peroxynitrite homeostasis using ROS/RNS conditional proteomics

**DOI:** 10.1101/2024.09.27.615315

**Authors:** Hao Zhu, Hiroaki Uno, Kyoichi Matsuba, Itaru Hamachi

## Abstract

Peroxynitrite (ONOO^−^), the product of the diffusion-controlled reaction of superoxide (O_2_^•−^) with nitric oxide (NO^•^), plays a crucial role in oxidative and nitrative stress and modulates key physiological processes such as redox signaling. While biological ONOO^−^ is conventionally analyzed using 3-nitrotyrosine antibodies and fluorescent sensors, such probes lack specificity and sensitivity, making high-throughput and comprehensive profiling of ONOO^−^-associated proteins challenging. In this study, we used a conditional proteomics approach to investigate ONOO^−^ homeostasis by identifying its protein neighbors in cells. We developed <u>P</u>er<u>o</u>xynitrite-<u>r</u>esponsive <u>p</u>rotein <u>L</u>abeling reagents (**Porp-L**) and, for the first time, discovered 2,6-dichlorophenol as an ideal moiety that can be selectively and rapidly activated by ONOO^−^ for labeling of proximal proteins. The reaction of **Porp-L** with ONOO^−^ generated several short-lived reactive intermediates that can modify Tyr, His, and Lys residues on the protein surface. We have demonstrated the **Porp-L**-based conditional proteomics in immune-stimulated macrophages, which indeed identified proteins known to be involved in the generation and modification of ONOO^−^ and revealed the endoplasmic reticulum (ER) as a ONOO^−^ hot spot. Moreover, we discovered a previously unknown role for Ero1a, an ER-resident protein, in the formation of ONOO^−^. Overall, **Porp-L** represent a promising research tool for advancing our understanding of the biological roles of ONOO^−^.

## Introduction

Reactive oxygen/nitrogen species (ROS/RNS) are a family of molecules continuously produced in living systems and involved in both physiology and pathology.^1, 2, 3^ Peroxynitrite (ONOO^−^) is a critical ROS/RNS family member that contributes to oxidative and nitrative stress in cells. It is produced by the non-enzymatic reaction of two primary ROS/RNS— superoxide (O_2_^•−^) and nitric oxide (NO^•^)—at a nearly diffusion-controlled rate (∼10^10^ M^−1^s^−1^)^4^ and outcompetes superoxide dismutase for the consumption of O_2_^•−^.^5^ The formation of ONOO^−^ is thus associated with the biogenesis of its parental species. While NO^•^ sources are restricted to nitric oxide synthases (NOSs),^6^ O_2_^•−^ arises from various cellular processes, such as NADPH oxidase (NOX) activation,^7^ mitochondrial electron leakage,^8^ and protein (mis)folding in the endoplasmic reticulum (ER).^9^ ONOO^−^ and the radicals derived from its secondary reactions readily cause oxidation and nitration of several protein targets, including transition metal centers (e.g., heme, [4Fe–4S] clusters), active cysteine (including thiolate ligands in zinc fingers), and tyrosine (Fig. 1a).^5, 10^ Such modifications can significantly alter protein structure and function, contributing to the pathogenesis of numerous maladies, such as inflammation,^11^ myocardial infarction,^12^ vascular diseases,^13^ and neurodegenerative disorders.^14^ Meanwhile, ONOO^−^ also acts as a modulator in several physiological events, including redox signaling,^15, 16^ platelet activation,^17^ and heart development.^18^ Despite having broad and important roles in health and disease, research on biological ONOO^−^ has lagged far behind that on other members of the ROS/RNS family.

**Figure 1.**
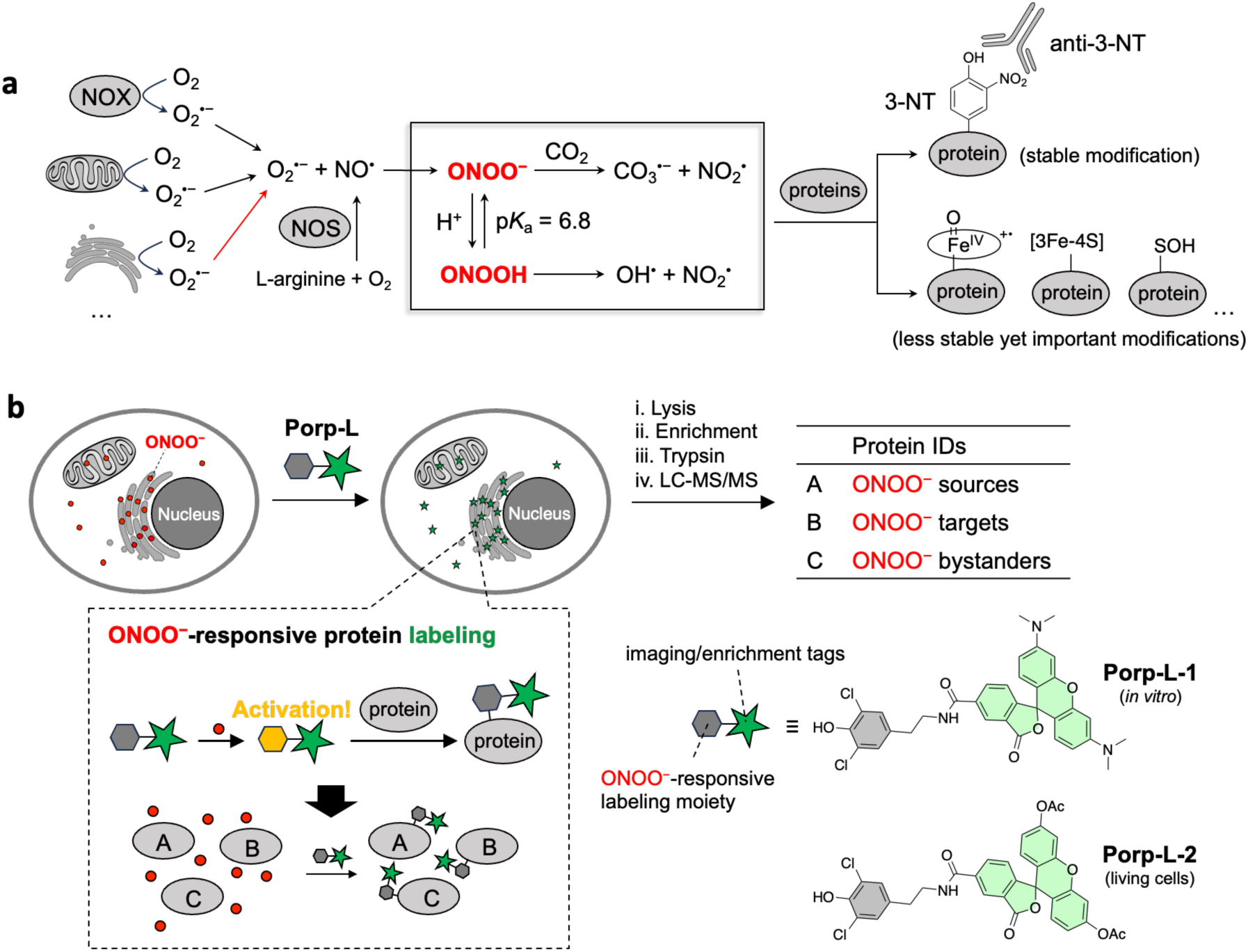
(a) Schematic illustration of ONOO^−^ biochemistry. NOX: NADPH oxidase. NOS: nitric oxide synthase. 3-NT: 3-nitrotyrosine. While NOX and mitochondrion have been the most extensively studied O_2_^·–^ sources, this work discovered Ero1a as an ER-resident O_2_^·–^ source for the formation of ONOO^−^ (red arrow). (b) Schematic illustration of **Porp-L**-based ROS/RNS conditional proteomics. The ONOO^−^ bystanders are not directly relevant to the generation and modification of ONOO^−^, while their identification also cues the localization of ONOO^−^.

Owing to the reactive and transient nature of ONOO^−^, its formation in biological systems has traditionally been assessed by analyzing its stable footprint, protein 3-nitrotyrosine (3-NT).^19, 20^ Antibodies that recognize 3-NT have proved to be particularly useful for identifying protein nitration in many disease-affected tissues.^21, 22, 23^ However, this method overlooks other, less stable yet important modifications induced by ONOO^−^, e.g., oxidation of transition metal centers and active cysteine, which determine the critical role of ONOO^−^ in enzyme inactivation. Additionally, 3-NT can be formed via ONOO^−^-independent pathways.^24^ More recently, the development of fluorescent probes has enabled direct and sensitive measurement of biological ONOO^−^ in living systems.^25, 26, 27, 28, 29^ To investigate the potential role of a hypothetical protein involved in ONOO^−^ homeostasis, such probes can be used to measure the change in ONOO^−^ concentration after genetic deletion of the protein.^30^ However, this method requires prior knowledge to firstly hypothesize a candidate protein and is inefficient for high-throughput and hypothesis-free discovery of multiple new proteins associated with ONOO^−^.

Herein, we report a new approach to detailing ONOO^−^ homeostasis, which utilizes ROS/RNS conditional proteomics to identify the protein neighbors of ONOO^−^ in living cells (Fig. 1b).^31, 32, 33^ This method relies on newly developed <u>P</u>er<u>o</u>xynitrite-responsive <u>p</u>rotein <u>L</u>abeling reagents (**Porp-L**) that can be selectively and quickly activated by ONOO^−^ and then transformed into highly reactive intermediates that immediately tag surrounding proteins. After cell lysis, the tagged proteins can be harvested via the enrichment handle in **Porp-L** before being digested with trypsin and analyzed using mass spectrometry (MS)-based proteomics. The identified ONOO^−^-surrounding proteins are strongly associated with the homeostasis and biochemistry of ONOO^−^, such as generation sources and modification targets. Moreover, proteins without a previously suggested role in ONOO^−^ homeostasis can be efficiently discovered using this method. In this study, we used **Porp-L**-based conditional proteomics in immune-stimulated macrophages to identify the ER as a hot spot of ONOO^−^ production and newly discover the involvement of Ero1a in ONOO^−^ formation.

## Results

### 1. Design of Porp-L

For conditional proteomics, a chemical probe is needed that can sense a particular target and enhance/trigger its reactivity to form covalent bonds with neighboring proteins.^31, 32, 33, 34, 35, 36, 37^ To date, there have been no reports on such probes for ONOO^−^, which should meet the following criteria: (1) high selectivity for ONOO^−^ over other ROS/RNS; (2) high sensitivity to ONOO^−^, enabling detection of low levels of endogenous ONOO^−^ in living systems; (3) fast kinetics of activation by ONOO^−^ and of subsequent protein labeling, which can minimize the lag between the initial ONOO^−^ encounter and final protein labeling and thus enable accurate analysis of ONOO^−^ neighbors in close proximity. Our design of **Porp-L** was inspired by the chemistry and biochemistry of ONOO^−^. It is known that ONOO^−^ reacts with phenol to produce nitrated and hydroxylated products.^38, 39^ We hypothesized that the intermediates in the reaction of ONOO^−^ with phenol could be exploited for protein labeling. For example, the phenoxyl radical, a well-established reactive species for proximity-dependent protein labeling,^40, 41, 42^ has been identified as a critical intermediate in 3-NT formation.^43^ To test this idea, we designed and synthesized a series of phenol derivatives (**1**–**4**) with various substituents in the *ortho* position(s) (Fig. 2a). Such substitutions were expected to modulate the reactivity of the phenol moiety with ONOO^−^ and also control the protein-reactivity of activated intermediates.

**Figure 2.**
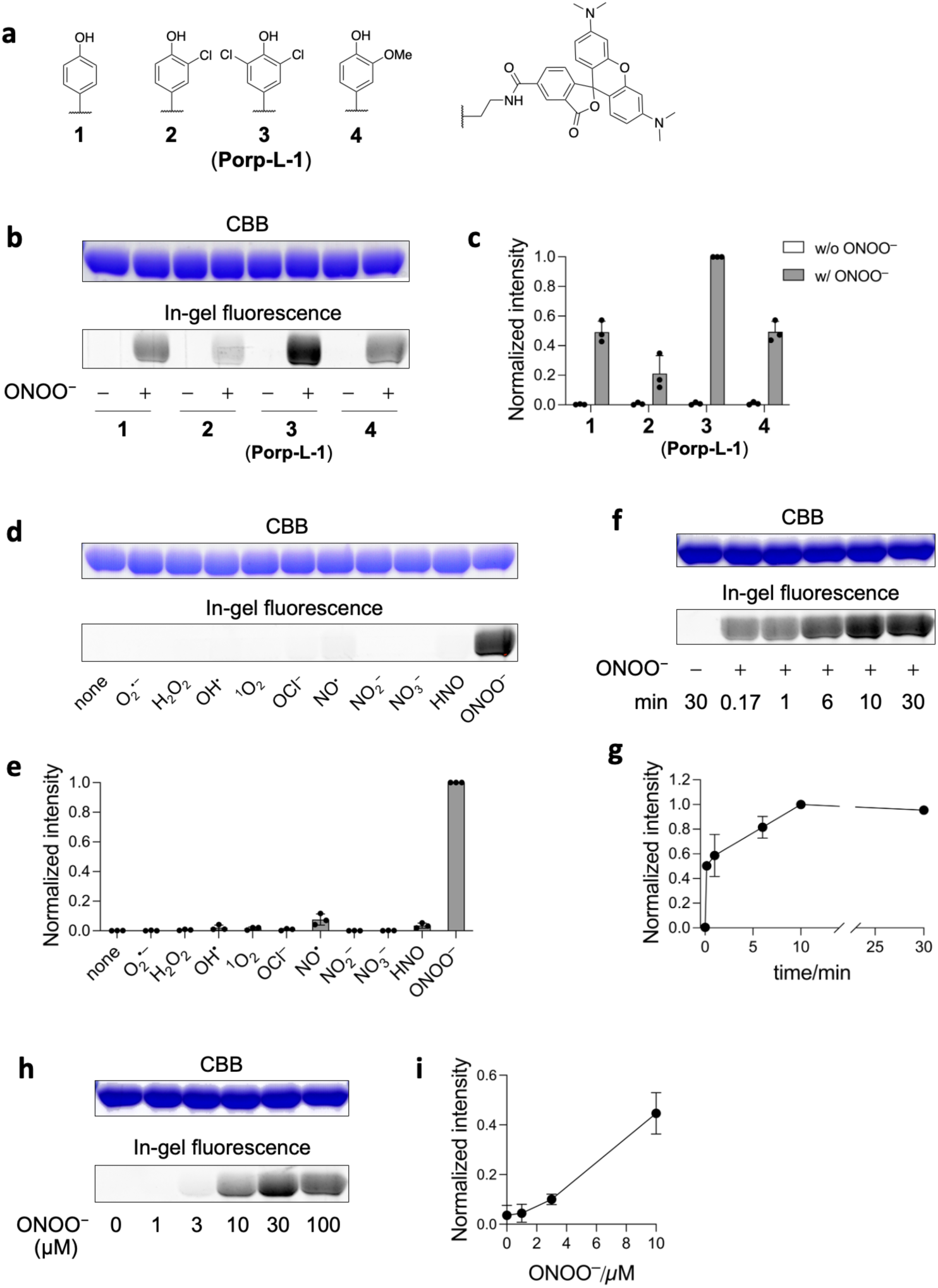
Characterization of ONOO^−^-responsive protein labeling *in vitro*. ONOO^−^ was added to a solution of BSA (10 µM) and probes **1-4** (10 µM) in phosphate buffer (0.1 M, pH 7.4). The labeling reaction was carried out at 37 °C. a) Chemical structures of probes **1-4**. b) Comparison of the protein labeling efficiency of probes **1-4** upon activation by ONOO^−^. ONOO^−^: 100 µM, 30 min. c) Quantification of the labeling in b). The in-gel fluorescence intensities were extracted and normalized relative to the ONOO^−^-induced labeling by **3**. d) ROS/RNS selectivity of **Porp-L-1**. ROS/RNS: 100 µM, 30 min. e) Quantification of the labeling in d). The intensities were normalized relative to the ONOO^−^-induced labeling. f) Time course of the ONOO^−^-induced labeling by **Porp-L-1**. ONOO^−^: 100 µM. g) Quantification of the labeling in f). The intensities were normalized relative to the labeling at 10 min. h) ONOO^−^ concentration-dependent protein labeling by **Porp-L-1**. ONOO^−^: 0-100 µM, 10 min. i) Quantification of the labeling in h). The intensities were normalized relative to the labeling at 30 µM of ONOO^−^. CBB: Coomassie brilliant blue. Error bars indicate s.d., n = 3.

### 2. Characterization of ONOO^−^-responsive protein labeling *in vitro*

The protein labeling propensity of probes **1**–**4** in response to ONOO^−^ was first examined using bovine serum albumin (BSA) as a model protein *in vitro*. Sodium dodecyl sulfate– polyacrylamide gel electrophoresis (SDS–PAGE) followed by in-gel fluorescence imaging showed that ONOO^−^-induced BSA labeling by **1**–**4** indeed occurred, and the efficiency was ranked in the following order: **3** > **1** ≈ **4** > **2** (Fig. 2b, c). This could be partially explained by the redox potentials and p*K*_a_ values of **1**–**4** (Table S1). Monochlorophenol (**2**) is relatively difficult to oxidize because of its high redox potential (0.93 V). Given the low p*K*_a_ (6.80) of dichlorophenol (**3**), it mainly exists as a phenolate anion at neutral pH, which may favor its oxidation by ONOO^−^. The ROS/RNS selectivities of **1**–**4** were then tested. Compared with **1** and **4**, probes **2** and **3** displayed superior selectivity for ONOO^−^ over a wide range of ROS and RNS, including O_2_^•−^ and NO^•^ (the parent molecules of ONOO^−^), H_2_O_2_ (one of the main cellular ROS), OH^•^ (derived from secondary reactions of ONOO^−^,^44^ among other sources), HNO (the reduction product of NO^•^)^45^, and other relevant reactive species (Fig. 2d, e and Fig. S1). On the basis of its efficient and selective activation by ONOO^−^, we further characterized probe **3**, which is hereafter named **Porp-L-1**.

We investigated the ONOO^−^-responsive protein labeling capacity of **Porp-L-1**. Kinetic analysis showed that the labeling reaction began immediately (≤10 s) following ONOO^−^ addition to a mixture of BSA and **Porp-L-1** and reached a plateau within 10 min (Fig. 2f, g). These observations are consistent with the short half-life (∼1 s) of ONOO^−^ at physiological pH^46^ and highlight the fast kinetics of both **Porp-L-1** activation by ONOO^−^ and subsequent protein labeling. Further BSA labeling analyses revealed that **Porp-L-1** is sensitive to ONOO^−^ at apparent concentrations as low as 3 µM (Fig. 2h, i), suggesting the potential of **Porp-L** to detect endogenous ONOO^−^ (produced at a rate of 50–100 µM min^−1^ in immune-stimulated macrophages^28^). **Porp-L-1** efficiently labeled protein in the presence of ONOO^−^ over a wide pH range, from pH 4 to 8 (Fig. S2), indicating the potential for **Porp-L** to function in diverse subcellular environments.

### 3. Activation and labeling mechanisms of Porp-L

We next attempted to investigate the ONOO^−^-induced activation and protein labeling mechanisms of **Porp-L**. A model compound (**Porp-L-M**) was used to simplify the analysis.

First, the reactivity of **Porp-L-M** toward various ROS/RNS was evaluated using high-performance liquid chromatography (HPLC) (Fig. 3a and Fig. S3). The intensity of the **Porp-L-M** elution peak decreased by approximately 25% in the presence of 1 equiv ONOO^−^, whereas it remained minimally altered after incubation with other ROS/RNS, including O_2_^•−^, H_2_O_2_, OH^•^, and NO^•^. These findings are consistent with the high selectivity of **Porp-L-1** for ONOO^−^ in BSA labeling (Fig. 2d, e). Subsequently, products of the reactions between **Porp-L-M** and ONOO^−^ in the presence and absence of a few of amino acids were analyzed using liquid chromatography–mass spectrometry (LC-MS) (Fig. S4). A dimer of **Porp-L-M**, (**Porp-L-M)_2_**, was detected following the reaction of **Porp-L-M** with ONOO^−^ in the absence of amino acids (Fig. 3b, c), and its structure was clearly characterized using proton nuclear magnetic resonance (^1^H-NMR) after purification (see Supporting Information). The dimer formation strongly suggests the generation of a dichlorophenoxyl radical (Intermediate 1) by one-electron oxidation of **Porp-L-M** (Fig. 3d). Consistent with a radical coupling reaction, an adduct of **Porp-L-M** with Tyr (i.e., Tyr labeling) was detected after incubation of **Porp-L-M** with Tyr and ONOO^−^. Moreover, we also identified conjugates of **Porp-L-M** with Cys, Lys, and His in the presence of ONOO^−^, indicating that ONOO^−^ triggers generation of other reactive intermediates likely derived from Intermediate 1. It was reported that disproportionation of phenoxyl radicals can produce a quinone methide^47^ (Intermediate 2), which enables to react with Cys, Lys, and His^33, 48^ (Cys1, Lys1, and His1 labeling). Two-electron oxidation of **Porp-L-M** can yield a positively charged dienone (Intermediate 3), which may conjugate with Cys (as detected) or react with a water molecule to generate a quinone^49, 50^ (Intermediate 4). In strong support of this potential intermediate, we identified adducts of Intermediate 4 with Lys and His^51^ (Lys2 and His2 labeling). The adduct of Cys with Intermediate 4 was not detected, likely because Cys outcompetes water in the reaction with Intermediate 3. In contrast, no conjugates of **Porp-L-M** with Ala, which lacks a nucleophilic side chain, were detected even in the presence of ONOO^−^. We also confirmed that glutathione (GSH), an abundant cellular nucleophile, also trapped Intermediates 2 and 3 of **Porp-L-M**, similar to Cys. Although GSH may therefore compromise the protein labeling efficiency of **Porp-L** in living systems, reaction of the intermediates with GSH can limit their diffusion and thus aid the spatial resolution of **Porp-L**-based labeling.

**Figure 3.**
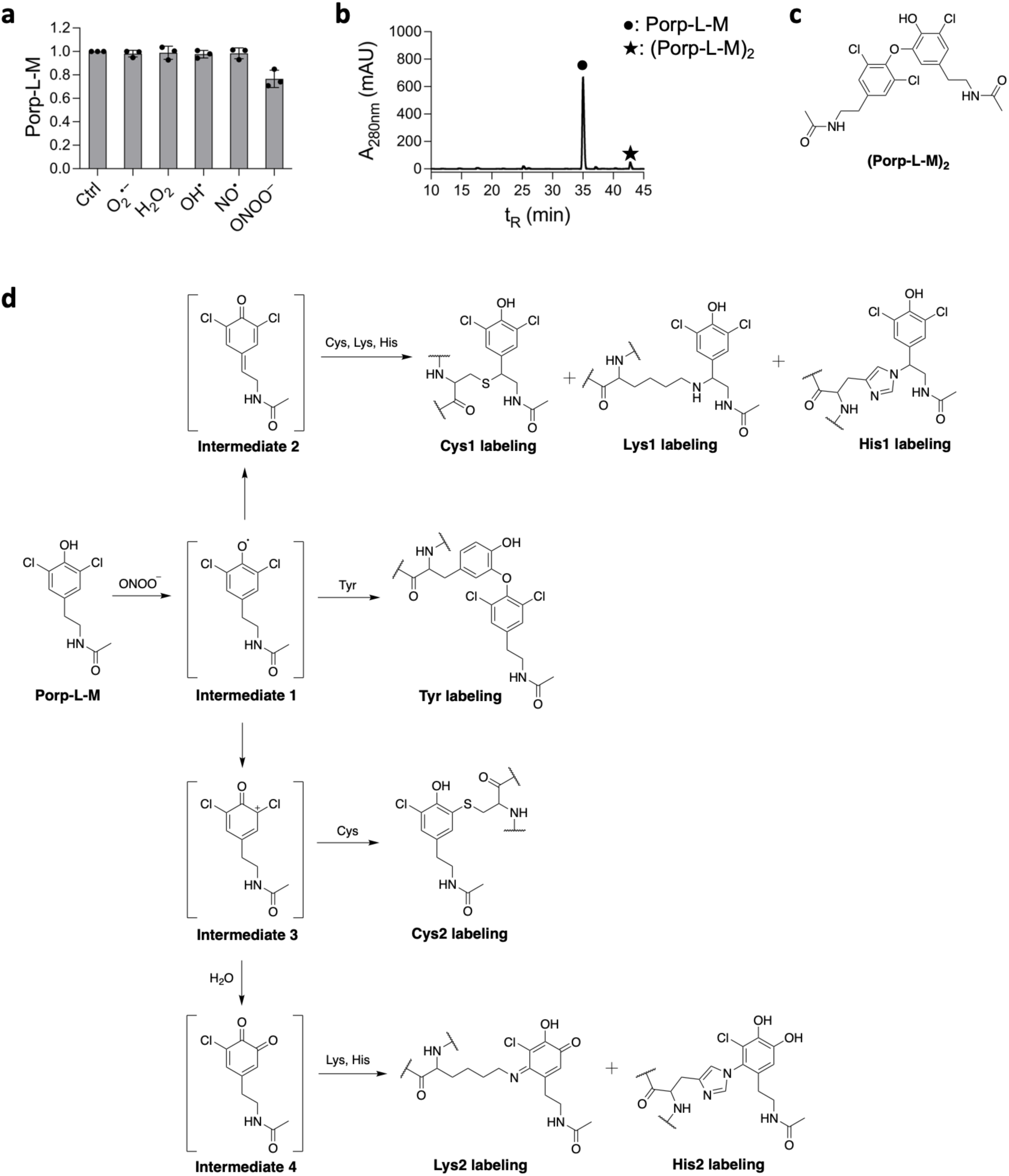
Investigating the ONOO^−^ activation and protein labeling mechanisms of **Porp-L**. a) Evaluation of the reactivity of **Porp-L-M** towards ROS/RNS. **Porp-L-M** (1 mM) was incubated with the indicated ROS/RNS (1 mM) in phosphate buffer (0.1 M, pH 7.4) at 37 °C for 30 min. The abundance of **Porp-L-M** was quantified according to the HPLC peak area and normalized relative to the control without ROS/RNS. Error bars indicate s.d. n = 3. b) LC-MS analysis of the reaction mixture of **Porp-L-M** with ONOO^−^. **Porp-L-M** (1 mM) was incubated with ONOO^−^ (1 mM) in phosphate buffer (0.1 M, pH 7.4) at 37 °C for 30 min. c) Structure of (**Porp-L-M)_2_**. d) Scheme of **Porp-L-M** reacting with ONOO^−^ and amino acids. The conjugates of **Porp-L-M** with amino acids were identified by LC-MS (Fig. S4).

Using the identified products of reaction with amino acids, we characterized the **Porp-L**-derived modification sites on BSA using liquid chromatography–tandem mass spectrometry (LC–MS/MS). After reaction of **Porp-L-M** with BSA in the presence of ONOO^−^, the protein was digested using trypsin and the resulting peptides were mapped using LC–MS/MS (Fig. 4a). The peptide hits were strictly defined by analyzing the characteristic chlorine isotope patterns of the diagnostic fragment ions (M1 and M2) of **Porp-L-M** and/or the peptide fragment ions (Fig. 4b, c and Fig. S5). In total, two Tyr residues (by Tyr labeling), two Lys residues (by Lys1 labeling), and six His residues (by His1 and His2 labeling) were modified by **Porp-L-M** (Fig. 4d and Table S2). All modified residues were found to be highly exposed on the protein surface (Fig. 4e), and such modifications were not observed in the absence of ONOO^−^. Collectively, these results show that **Porp-L** can be activated by ONOO^−^ to generate several reactive intermediates that can broadly modify amino acids exposed on a protein surface.

**Figure 4.**
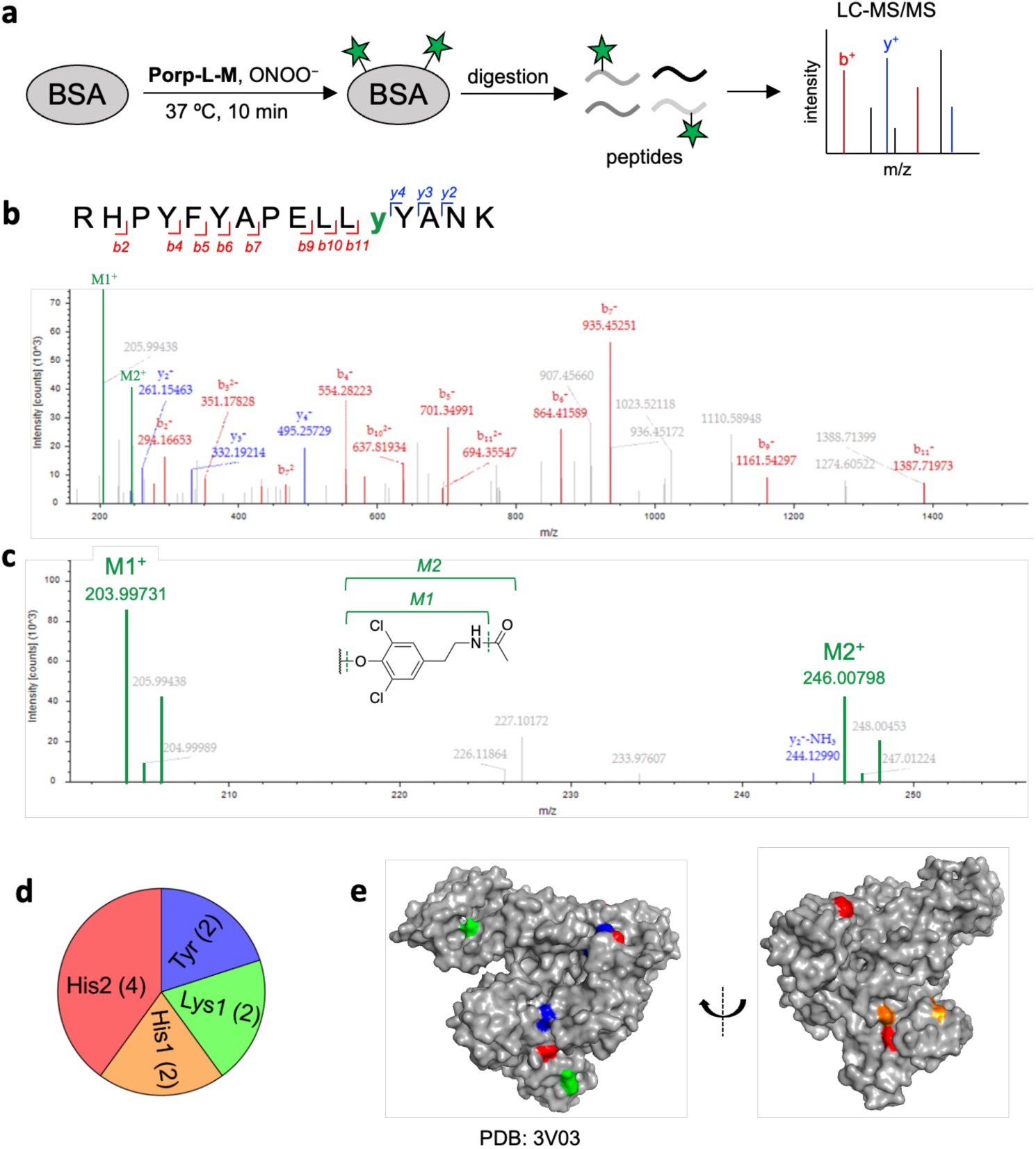
Analysis of **Porp-L**-labeled amino acid residues on BSA. ONOO^−^ (5 mM) was added to a solution of BSA (100 µM) and **Porp-L-M** (1 mM) in phosphate buffer (0.1 M, pH 7.4). The labeling reaction was carried out at 37 °C for 10 min. a) Scheme of peptide mapping of **Porp-L-M**-modified BSA. b) MS/MS spectrum of one **Porp-L-M**-modified peptide. M1 and M2 denote the fragment ions derived from **Porp-L-M**. c) A zoom window of b) shows the characteristic chlorine isotope pattern of the M1 and M2 fragments. d) Pie chart of **Porp-L-M**-modified residues on BSA. e) Crystal structure of BSA visualized by PyMOL. **Porp-L-M**-modified residues were highlighted. Tyr: blue, Lys: green, His1: orange, His2, red.

### 4. Application of Porp-L in living cells

To test **Porp-L** in living cells, a cell-permeable probe, **Porp-L-2**, bearing fluorescein diacetate was synthesized. Using confocal laser scanning microscopy (CLSM), we first confirmed that **Porp-L-2** could rapidly permeate live RAW264.7 macrophages and achieve a broad intracellular distribution (Fig. 5a). SDS–PAGE followed by in-gel fluorescence imaging showed that **Porp-L-2** was stable in living cells and resulted in negligible protein labeling after 60 min incubation. When 3-(4-morpholinyl)sydnonimine (SIN-1), a synthetic ONOO^−^ generator, was added to the **Porp-L-2**-pretreated cells, significant protein labeling was observed (Fig. S6). The presence of multiple fluorescent protein bands over a wide molecular weight range was suggestive of promiscuous labeling by **Porp-L-2**.

**Figure 5.**
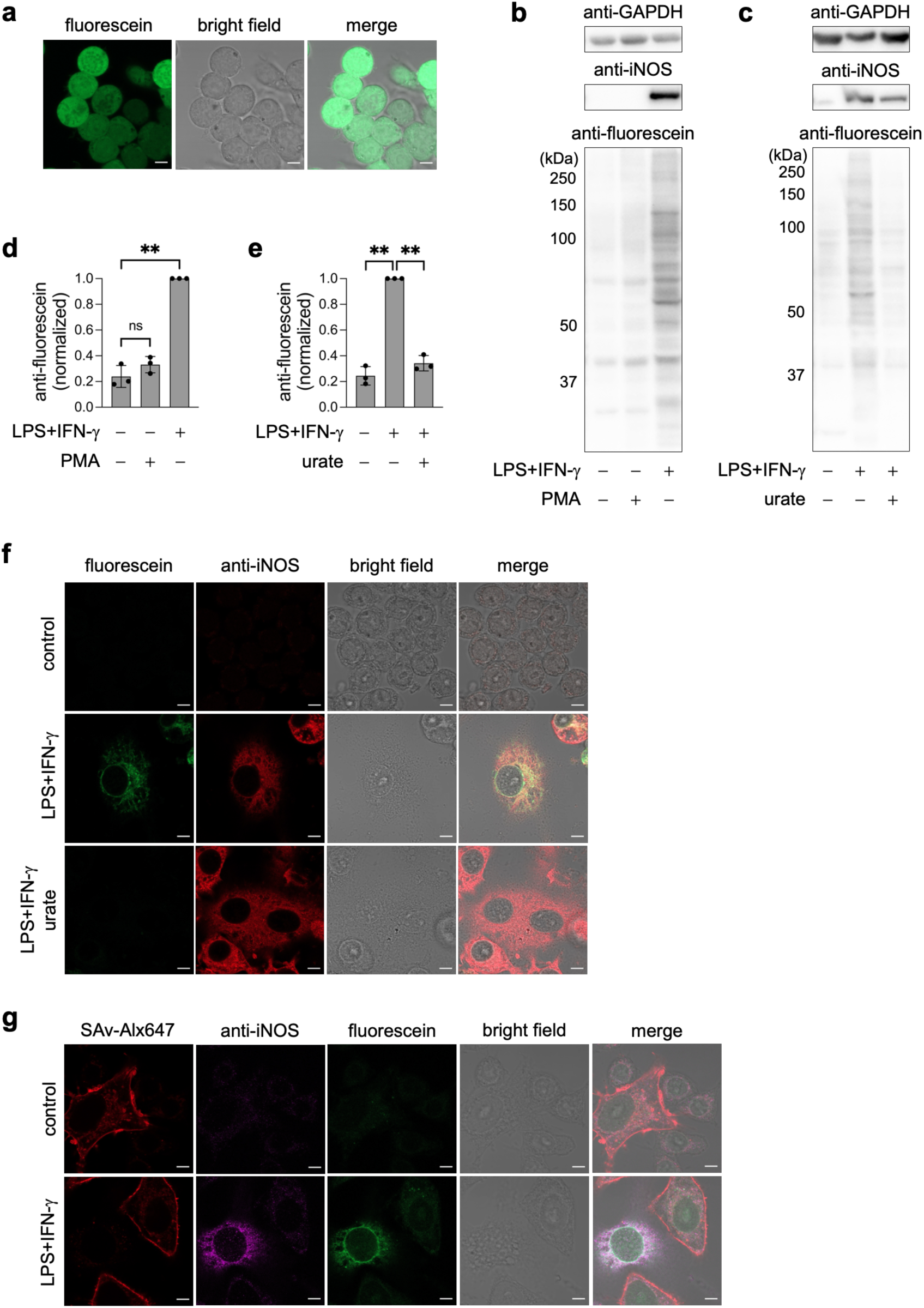
Application of **Porp-L** in living cells. a) Live-cell distribution of **Porp-L-2**. RAW264.7 macrophages were incubated with **Porp-L-2** (5 µM) at 37 °C for 5 min. b-c) Western blot analysis of protein labeling by **Porp-L-2** in response to endogenously produced ONOO^−^ upon immune stimulation. RAW264.7 macrophages were treated with PMA (0.2 µg/mL) for 0.5 h or LPS (0.5 µg/mL) and IFN-γ (7.5 ng/mL) for 24 h, followed by incubation with **Porp-L-2** (5 µM) for 1 h. Urate (1 mM) was added 1 h prior to the labeling by **Porp-L-2**. The protein labeling was analyzed by Western blot using an anti-fluorescein antibody. d-e) Quantification of the protein labeling by **Porp-L-2** in b-c). Error bars indicate s.d., n = 3. Statistical analyses were performed with a two-tailed Student’s t-test, ns: not statistically significant, \*\**P* < 0.01. f) CLSM imaging of the protein labeling by **Porp-L-2**. The cells were fixed by cold methanol. Scale bar: 5 µm. g) Selective labeling of ONOO^−^-producing cells in mixed cell culture. RAW264.7 macrophages were mixed with surface-biotinylated HeLa cells, followed by the LPS + IFN-γ stimulation and **Porp-L-2** labeling, as described above. The cells were fixed with cold methanol and HeLa cells were fluorescently marked by a SAv-Alx647 conjugate. Scale bar: 5 µm.

We then assessed the performance of **Porp-L-2** in response to endogenous ONOO^−^ in RAW264.7 macrophages upon immune stimulation. The cells were challenged for 24 h with the bacterial endotoxin lipopolysaccharide (LPS) and the pro-inflammatory cytokine interferon-ψ (IFN-ψ), inducing the upregulation of inducible NOS (iNOS) (Fig. 5b) and the production of endogenous ONOO^−^ (Fig. S7).^27, 52, 53^ **Porp-L-2** was added to the stimulated cells and incubated for 60 min. Western blot analysis revealed significant protein labeling by **Porp-L-2** in the stimulated cells (Fig. 5b, d). Preincubation with 1400w (a selective iNOS inhibitor)^54^ or urate (a ONOO^−^ scavenger)^55^ markedly attenuated the protein labeling (Fig. 5c, e, and Fig. S8), revealing that the **Porp-L-2** labeling was mainly induced by endogenous ONOO^−^. More interestingly, western blot using an anti-3-NT antibody could not clearly detect nitrated proteins under the identical stimulation condition using LPS and IFN-γ (Fig. S9), and obvious protein nitration was observed only when the cell lysates were treated with a millimolar concentration of ONOO^−^. This highlights the advantage of **Porp-L** for the direct and sensitive investigation of ONOO^−^ homeostasis in living cells. We also tested the behavior of **Porp-L-2** in RAW264.7 macrophages treated with phorbol myristate acetate (PMA) to activate NOX, which catalyzes the production of O_2_^•−^, H_2_O_2_, and OH^•^ but not ONOO^−^.^33, 56^ Consistent with its specificity for ONOO^−^, **Porp-L-2** resulted in negligible protein labeling in the PMA-treated cells (Fig. 5b, d). These results demonstrated the ability of **Porp-L-2** to label proteins in response to endogenous ONOO^−^ produced under immune stimulation and confirmed its high selectivity for ONOO^−^ in the complex ROS/RNS environment of living cells. The intracellular distribution of the fluorescein-tagged proteins by **Porp-L-2** was visualized using CLSM after cell fixation. In contrast to the nearly homogenous distribution of **Porp-L-2** in live cells (Fig. 5a), its labeling was limited to certain subcellular regions of the immune-stimulated macrophages, indicating a spatially heterogenous intracellular production of ONOO^−^ (Fig. 5f and Fig. S10). Additional immunostaining assays revealed that **Porp-L-2**-labeled proteins partially colocalized with iNOS, suggesting that some ONOO^−^ was likely formed close to where NO^•^ was synthesized. Consistent with the above western blot results (Fig. 5b and Fig. S8), control experiments involving non-stimulation of cells, iNOS inhibition, and ONOO^−^ scavenging resulted in negligible intracellular fluorescence.

We further tested **Porp-L-2** in a co-culture system to examine the intercellular diffusion of ONOO^−^. In contrast to that in RAW264.7 macrophages, LPS and IFN-γ could not induce the expression of iNOS and any increased protein labeling by **Porp-L-2** in HeLa cells (Fig. S11). A co-culture of RAW264.7 macrophages and HeLa cells were stimulated with LPS and IFN-γ and thereafter incubated with **Porp-L-2**. CLSM after cell fixation revealed marked protein labeling by **Porp-L-2** in RAW264.7 macrophages expressing iNOS, whereas minimal labeling was observed in the nearby HeLa cells (Fig. 5g and Fig. S12). This result suggested that ONOO^−^ is primarily localized within the cells where it is formed and undergoes minimal extracellular leakage, which is likely a consequence of its high reactivity with various biomolecules and short half-life.

### 5. ROS/RNS conditional proteomics

For in-depth analysis of the proteins associated with ONOO^−^ homeostasis, we carried out conditional proteomics to identify the **Porp-L-2**-labeled proteins (Fig. 6a). After stimulation with LPS and IFN-γ and labeling with **Porp-L-2**, the cells were lysed and the labeled proteins were harvested via immunoprecipitation with an anti-fluorescein antibody. Quantitative mass analysis was performed by label-free quantification (LFQ) against a non-stimulated control. A total of 243 proteins were detected, and 32 proteins that met the criteria of fold-change > 2 and *P* < 0.05 were regarded as hit proteins (Fig. 6b and Table S3).

**Figure 6.**
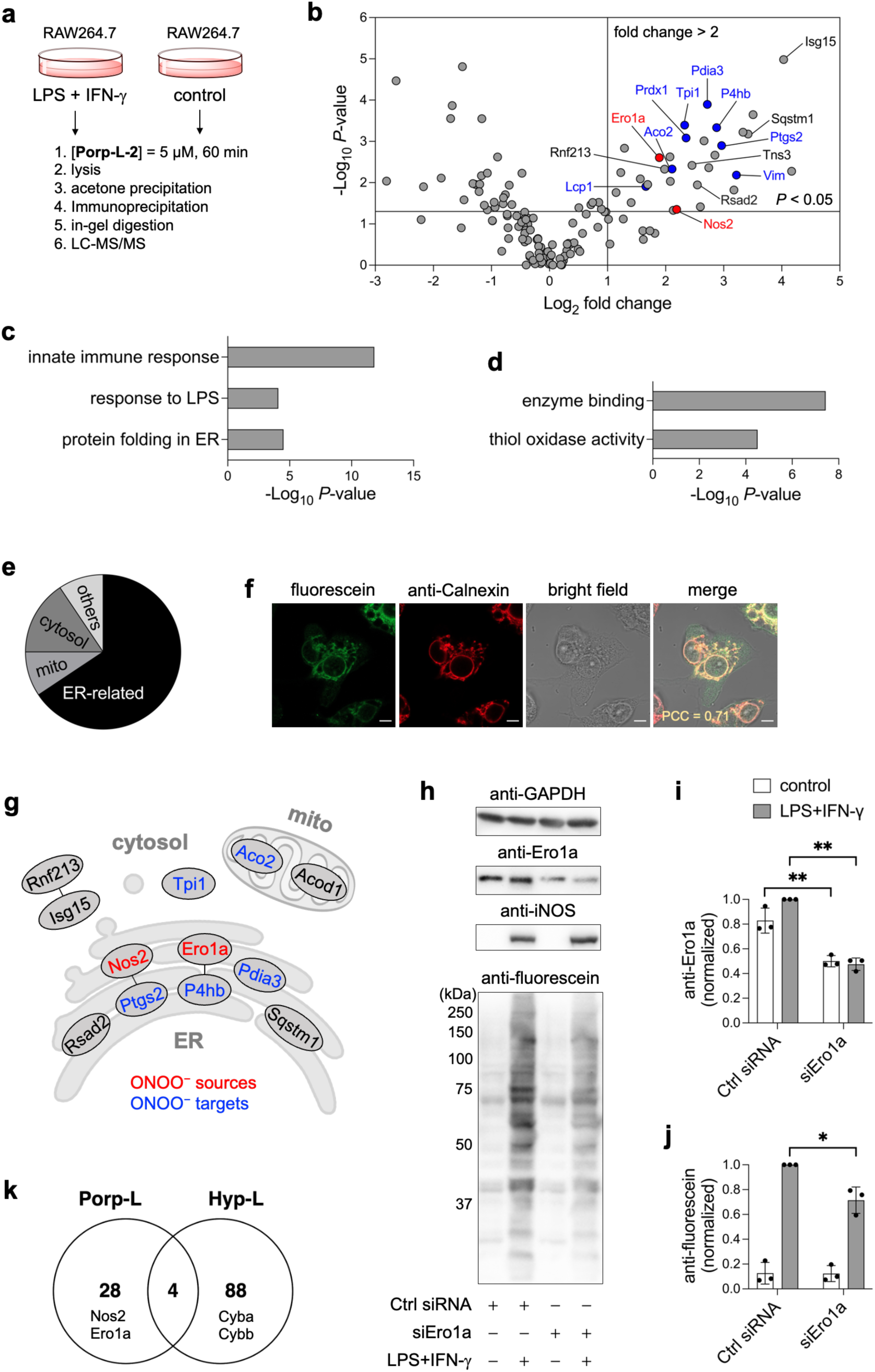
Profiling ONOO^−^-surrounding proteins by ROS/RNS proteomics. a) Workflow of the **Porp-L**-based ROS/RNS conditional proteomics. b) Volcano plot obtained by LFQ quantitative proteomics (LPS+IFN-γ/control) with three biological replicates. Red: ONOO^−^ sources, blue: ONOO^−^ targets. Biological Process (c) and Molecular Function (d) analysis of the hit proteins. e) Classification of the hit proteins based on subcellular localization. f) Co-localization of the **Porp-L** labeling with an ER marker (anti-calnexin). The cells were fixed with cold methanol. Scale bar: 5 µm. g) Mapping of the representative hit proteins based on their subcellular localization and association with ONOO^−^. The edges indicate physical protein interactions (https://string-db.org/). h) SiRNA knockdown of Ero1a. RAW264.7 macrophages were transfected with control or Ero1a-targeting siRNAs for 24 h, followed by the LPS + IFN-γ stimulation and **Porp-L-2** labeling. Quantification of the Ero1a expression (i) and the **Porp-L** labeling (j) in h). Error bars indicate s.d., n = 3. Statistical analyses were performed with a two-tailed Student’s t-test. \**P* < 0.05, \*\**P* < 0.01.

First, the hit proteins were globally analyzed using Gene Ontology (GO) and found to be significantly related to biological processes and molecular functions including innate immune response, protein folding in the ER, and thiol oxidase activity (Fig. 6c, d). Cellular component analysis assigned 21 hit proteins (66%) to the ER, three to the mitochondrion (9%), and five to the cytosol (16%) (Fig. 6e). Immunofluorescence staining and CLSM showed that proteins labeled with **Porp-L-2** indeed highly colocalized (Pearson’s correlation coefficient = 0.71) with calnexin, an ER marker (Fig. 6f and Fig. S13). The global proteomics and CLSM results strongly suggested that the endogenous ONOO^−^ induced by LPS and IFN-γ stimuli was mainly produced and/or accumulated in the ER. This agrees with our previous finding that immune-stimulated RAW264.7 macrophages have a high level of NO^•^ in the ER.^32^

Next, we individually annotated the hit proteins (Fig. 6b, g). As expected, Nos2 (iNOS), a central NO^•^ source in macrophages, was identified by **Porp-L**-based proteomics, a finding consistent with the colocalization of the **Porp-L-2** labeling and iNOS immunostaining described above (Fig. 5f). Several known ONOO^−^ targets were also found in the hit protein list, such as Aco2 (Fe–S cluster oxidation; mitochondrion), Tpi1 (Tyr nitration; cytosol), Ptgs2 (Tyr nitration; ER), and P4hb (Cys oxidation; ER). While other hit proteins, including Rsad2, Rnf213, Sqstm1, and Tns3, have not previously been identified as ONOO^−^ targets, we noticed that each of them contains a potentially reactive site that could be susceptible to attack by ONOO^−^, namely a [4Fe–4S] cluster (Rsad2), zinc finger (Rnf213 and Sqstm1), and a CXXXXXR motif (Tns3). S-nitrosylation (a hallmark modification by NO^•^) of Rnf213 was previously identified in the brains of tauopathy model mice with elevated oxidative and nitrative stress.^57^ Isg15, a Rnf213 interactor^58^ that can be S-nitrosylated for effective protein ISGylation during an immune response,^59^ was also identified using our **Porp-L**-based proteomics. Moreover, on the basis of the similar anionic structures of ONOO^−^ and phosphate, a previous report suggested that ONOO^−^ can enter the active site of a phosphatase possessing the CXXXXXR motif (which also appears in Tns3) and thereby oxidize the catalytic Cys.^60^

Among the proteins newly revealed by **Porp-L**-based proteomics, we focused on Ero1a, an ER-resident protein that is known to regulate oxidative protein folding by re-oxidizing reduced protein disulfide isomerase and to produce ROS by passing electrons to molecular oxygen.^9, 61^ To date, no reports have identified or even suggested an association between Ero1a and ONOO^−^. To validate the involvement of Ero1a in ONOO^−^ homeostasis, we carried out siRNA knockdown of Ero1a in RAW264.7 macrophages, which suppressed the expression of Ero1a without impacting that of iNOS (Fig. 6h, i). The western blot data clearly showed that the amount of protein labeled with **Porp-L-2** was substantially decreased (by approximately 30%) after the knockdown of Ero1a in the stimulated cells (Fig. 6h, j). Given its ability to generate ROS in the ER, it is reasonable to conclude that Ero1a may serve as one of O_2_^•−^ sources for the formation of ONOO^−^ rather than being simply a ONOO^−^ bystander. These experiments highlighted that **Porp-L**-based conditional proteomics can specifically and efficiently reveal the proteins associated with ONOO^−^ homeostasis in living cells.

Lastly, we sought to perform multiplexed conditional proteomics using different ROS/RNS-specific chemical probes. We previously developed an H_2_O_2_-responsive protein labeling reagent (**Hyp-L**) for H_2_O_2_-focused proteomics.^33^ Here we again conducted **Hyp-L**-based proteomics in PMA-activated RAW264.7 macrophages under the conditions for lysis, immunoprecipitation, and LC-MS/MS, identical with those of **Porp-L**. Using **Hyp-L**, 92 hit proteins were identified that met the criteria of fold-change > 2 and *P* < 0.05 (Fig. S14 and Table S4). Comparison of the ONOO^−^ proteome obtained using **Porp-L** with the H_2_O_2_ proteome obtained using **Hyp-L** revealed only four proteins in common (Fig. 6k). Importantly, key proteins relevant to ONOO^−^ and H_2_O_2_ homeostasis were exclusively targeted by **Porp-L** and **Hyp-L**, respectively. For instance, Nos2 and Ero1a were solely categorized into the **Porp-L** proteome, while Cyba and Cybb, the core subunits of NOX responsible for H_2_O_2_ production in PMA-activated macrophages, were exclusively found in the **Hyp-L** proteome. Thus, by using ROS/RNS-specific protein labeling reagents, conditional proteomics enables proteins associated with distinct ROS/RNS to be profiled and differentiated and has the potential to comprehensively characterize ROS/RNS homeostasis in multiplex assays.

## Discussion

We developed a ONOO^−^-focused chemical proteomics approach, which tags and identifies ONOO^−^-surrounding proteins that are relevant to the generation, modification, and localization of ONOO^−^ in biological systems. Our initial efforts focused on developing chemical reagents for ONOO^−^-responsive protein labeling (**Porp-L**), which has never been achieved before. For the first time, we identified 2,6-dichlorophenol as an effective ONOO^−^-responsive moiety that can be selectively activated by ONOO^−^ for efficient protein labeling. **Porp-L** exhibited rapid kinetics of both activation by ONOO^−^ and subsequent protein labeling, making it ideal for precisely tagging ONOO^−^-proximal proteins. Several short-lived, protein-reactive intermediates were derived from the reaction of **Porp-L** with ONOO^−^, including the phenoxyl radical, quinone methide, positively charged dienone, and quinone, which can broadly modify amino acid residues (Tyr, His, and Lys) on the protein surface.

When applied to living cells, **Porp-L** resulted in promiscuous protein labeling in areas of endogenous ONOO^−^ generation. Covalent protein tagging with **Porp-L** permanently stamped the transient ONOO^−^, which can be visualized using fluorescence microscopy after cell fixation. This highlights an advantage of **Porp-L** over current fluorescent sensors which may readily diffuse after reacting with ONOO^−^ and usually do not survive cell fixation. More importantly, following our conditional proteomics workflow, **Porp-L**-labeled proteins can be harvested and identified, which can efficiently reveal various proteins involved in ONOO^−^ homeostasis. Notably, **Porp-L** labeled several proteins having a transition metal center or an active cysteine, which are prevalent biological targets of ONOO^−^ but cannot be surveyed using traditional 3-NT-focused analysis. Consequently, **Porp-L** enables more sensitive and comprehensive analyses of ONOO^−^ than an anti-3-NT antibody (Fig. S9). Using **Porp-L**-based conditional proteomics and CLSM, we indeed demonstrated that endogenously produced ONOO^−^ was heterogeneously distributed in cells and condensed in the ER. Also, movement of ONOO^−^ between cells was shown to be minimal, which is consistent with its high reactivity and short lifetime. Furthermore, our approach not only identified the NO^•^-producing enzyme in macrophages and known protein targets of ONOO^−^ but also uncovered proteins potentially involved in ONOO^−^ generation and modification, which enabled us to discover a previously unknown role for Ero1a in ONOO^−^ generation.

We envision the widespread use of **Porp-L** to elucidate the roles of ONOO^−^ in various physiological and pathological processes, such as protein tyrosine phosphorylation, aging, inflammation, and neurodegeneration in cultured cells as well as more complex environments including primary cultures, tissues, and live animals. By combining **Porp-L** with other ROS/RNS-responsive protein labeling reagents that we have previously developed,^32, 33^ we aim to achieve a comprehensive understanding of ROS/RNS homeostasis, with detailed insights into specific ROS/RNS family members.

## Acknowledgments

We thank Edanz (https://jp.edanz.com/ac) for editing a draft of this manuscript. This work was supported by a Grant-in-Aid for Specially Promoted Research (JSPS KAKENHI Grant 23H05405) and the Japan Science and Technology Agency (JST) ERATO Grant JPMJER1802 to I.H., and a Grant-in-Aid for Early-Career Scientists (23K13855) to H.Z.

## Author Contributions

H.Z. and I.H. conceived the project and designed the experiments. H.U. synthesized the compounds and carried out the in-tube experiments. H.Z. conducted the in-cell experiments. K.M. prepared the samples for proteomics. H.Z. and I.H. wrote the manuscript with inputs from all the authors.

## Competing Interests statement

The authors declare no competing interests.

## Notes

### Competing Interest Statement

The authors have declared no competing interest.

## References

1. Winterbourn, C. C. Reconciling the chemistry and biology of reactive oxygen species. Nat. Chem. Biol. 4, 278–286 (2008).

2. Dickinson, B. C., Chang, C. J. Chemistry and biology of reactive oxygen species in signaling or stress responses. Nat. Chem. Biol. 7, 504–511 (2011).

3. Sies, H. et al. Defining roles of specific reactive oxygen species (ROS) in cell biology and physiology. Nat. Rev. Mol. Cell Biol., 23, 499–515 (2022).

4. Huie, R. E., Padmaja, S. The reaction of NO with superoxide. Free Radic. Res. Commun. 18, 195–199 (1993).

5. Pacher, P., Beckman, J. S., Liaudet, L. Nitric oxide and peroxynitrite in health and disease. Physiol. Rev. 87, 315–424 (2007).

6. Förstermann, U., Sessa, W. C. Nitric oxide synthases: regulation and function. Eur. Heart J. 33, 829–837 (2012).

7. Bedard, K., Krause, K.-H. The NOX family of ROS-generating NADPH oxidases: physiology and pathophysiology. Physiol. Rev. 87, 245–313 (2007).

8. Nolfi-Donegan, D., Braganza, A., Shiva, S. Mitochondrial electron transport chain: oxidative phosphorylation, oxidant production, and methods of measurement. Redox Biol. 37, 101674–101674 (2020).

9. Tu, B. P., Weissman, J. S. Oxidative protein folding in eukaryotes: mechanisms and consequences. J. Cell Biol. 164, 341–346 (2004).

10. Ferrer-Sueta, G. et al. Biochemistry of peroxynitrite and protein tyrosine nitration. Chem. Rev. 118, 1338–1408 (2018).

11. van der Veen, R. C., Hinton, D. R., Incardonna, F., Hofman, F. M. Extensive peroxynitrite activity during progressive stages of central nervous system inflammation. J. Neuroimmunol. 77, 1–7 (1997).

12. Ronson, R. S., Nakamura, M., Vinten-Johansen, J. The cardiovascular effects and implications of peroxynitrite. Cardiovasc. Res. 44, 47–59 (1999).

13. Förstermann, U., Münzel, T. Endothelial nitric oxide synthase in vascular disease. Circulation 113, 1708–1714 (2006).

14. Torreilles, F., Salman-Tabcheh, S. d., Guérin, M.-C., Torreilles, J. Neurodegenerative disorders: the role of peroxynitrite. Brain Res. Rev. 30, 153–163 (1999).

15. Sawa, T. et al. Protein S-guanylation by the biological signal 8-nitroguanosine 3′,5′-cyclic monophosphate. Nat. Chem. Biol. 3, 727–735 (2007).

16. Monteiro, H. P., Arai, R. J., Travassos, L. R. Protein tyrosine phosphorylation and protein tyrosine nitration in redox signaling. Antioxid. Redox Signal. 10, 843–890 (2008).

17. Naseem, K. M. et al. The nitration of platelet cytosolic proteins during agonist-induced activation of platelets. FEBS Lett. 473, 119–122 (2000).

18. Viera, L. et al. Temporal patterns of tyrosine nitration in embryo heart development. Free Radic. Biol.Med. 55, 101–108 (2013).

19. Batthyány, C. et al. Tyrosine-nitrated proteins: proteomic and bioanalytical aspects. Antioxid. Redox Signal. 26, 313–328 (2017).

20. Viera, L., Ye, Y. Z., Estévez, A. G., Beckman, J. S. Immunohistochemical methods to detect nitrotyrosine. Methods Enzymol. 301, 373–381 (1999).

21. Beckmann, J. S. et al. Extensive nitration of protein tyrosines in human atherosclerosis detected by immunohistochemistry. Biol. Chem. Hoppe-Seyler 375, 81–88 (1994).

22. Smith, M. A., Harris, P. L. R., Sayre, L. M., Beckman, J. S., Perry, G. Widespread peroxynitrite-mediated damage in Alzheimer’s disease. J. Neurosci. 17, 2653–2657 (1997).

23. Ischiropoulos, H. Biological tyrosine nitration: a pathophysiological function of nitric oxide and reactive oxygen species. Arch. Biochem. Biophys. 356, 1–11 (1998).

24. Mohiuddin, I. et al. Nitrotyrosine and chlorotyrosine: clinical significance and biological functions in the vascular system. J. Surg. Res. 133, 143–149 (2006).

25. Ueno, T., Urano, Y., Kojima, H., Nagano, T. Mechanism-based molecular design of highly selective fluorescence probes for nitrative stress. J. Am. Chem. Soc. 128, 10640–10641 (2006).

26. Yang, D., Wang, H.-L., Sun, Z.-N., Chung, N.-W., Shen, J.-G. A highly selective fluorescent probe for the detection and imaging of peroxynitrite in living cells. J. Am. Chem. Soc. 128, 6004–6005 (2006).

27. Peng, T. et al. Molecular imaging of peroxynitrite with HKGreen-4 in live cells and tissues. J. Am. Chem. Soc. 136, 11728–11734 (2014).

28. Rios, N. et al. Sensitive detection and estimation of cell-derived peroxynitrite fluxes using fluorescein-boronate. Free Radic. Biol. Med. 101, 284–295 (2016).

29. Chen, Z., Zhang, S., Li, X., Ai, H.-W. A high-performance genetically encoded fluorescent biosensor for imaging physiological peroxynitrite. Cell Chem. Biol. 28, 1542–1553 (2021).

30. Kameritsch, P. et al. The mitochondrial thioredoxin reductase system (TrxR2) in vascular endothelium controls peroxynitrite levels and tissue integrity. Proc. Nat. Acad. Sci. 118, e1921828118 (2021).

31. Miki, T. et al. A conditional proteomics approach to identify proteins involved in zinc homeostasis. Nat. Methods 13, 931–937 (2016).

32. Nishikawa, Y. et al. Development of a nitric oxide-responsive labeling reagent for proteome analysis of live cells. ACS Chem. Biol. 14, 397–404 (2019).

33. Zhu, H. et al. Imaging and profiling of proteins under oxidative conditions in cells and tissues by hydrogen-peroxide-responsive labeling. J. Am. Chem. Soc. 142, 15711–15721 (2020).

34. Lee, S. et al. Activity-based sensing with a metal-directed acyl imidazole strategy reveals cell type-dependent pools of labile brain copper. J. Am. Chem. Soc. 142, 14993–15003 (2020).

35. Iwashita, H., Castillo, E., Messina, M. S., Swanson, R. A. Chang, C. J. A tandem activity-based sensing and labeling strategy enables imaging of transcellular hydrogen peroxide signaling. Proc. Nat. Acad. Sci. 118, e2018513118 (2021).

36. Pezacki, A. T. et al. Oxidation state-specific fluorescent copper sensors reveal oncogene-driven redox changes that regulate labile copper(II) pools. Proc. Nat. Acad. Sci. 119, e2202736119 (2022).

37. Cheng, R. et al. Protein-labeling reagents selectively activated by copper(I). ACS Chem. Biol. 19, 1222–1228 (2024).

38. van der Vliet, A., O’Neill, C. A., Halliwell, B., Cross, C. E., Kaur, H. Aromatic hydroxylation and nitration of phenylalanine and tyrosine by peroxynitrite. FEBS Lett. 339, 89–92 (1994).

39. Nonoyama, N., Chiba, K., Hisatome, K., Suzuki, H., Shintani, F. Nitration and hydroxylation of substituted phenols by peroxynitrite. Tetrahedron Lett. 40, 6933–6937 (1999).

40. Hyun-Woo, R. et al. Proteomic mapping of mitochondria in living cells via spatially restricted enzymatic tagging. Science 339, 1328–1331 (2013).

41. Loh, K. H. et al. Proteomic analysis of unbounded cellular compartments: synaptic clefts. Cell 166, 1295–1307 (2016).

42. Oslund, R. C. et al. Detection of cell–cell interactions via photocatalytic cell tagging. Nat. Chem. Biol., 18, 850–858 (2022).

43. Gunaydin, H., Houk, K. N. Mechanisms of peroxynitrite-mediated nitration of tyrosine. Chem. Res. Toxicol. 22, 894–898 (2009).

44. Szabó, C., Ischiropoulos, H., Radi, R. Peroxynitrite: biochemistry, pathophysiology and development of therapeutics. Nat. Rev. Drug Discov. 6, 662–680 (2007).

45. Suarez, S. A. et al. HNO is produced by the reaction of NO with thiols. J. Am. Chem. Soc. 139, 14483–14487 (2017).

46. Hughes, M. N. Chemistry of nitric oxide and related species. Methods Enzymol. 436, 3–19 (2008).

47. Stebbins, R., Sicilio, F. The kinetics of disproportionation of the 2,6-di-t-butyl-4-methyl phenoxy radical. Tetrahedron 26, 291–297 (1970).

48. Bolton, J. L., Turnipseed, S. B., Thompson, J. A. Influence of quinone methide reactivity on the alkylation of thiol and amino groups in proteins: studies utilizing amino acid and peptide models. Chem. Biol. Interact. 107, 185–200 (1997).

49. Osborne, R. L., Coggins, M. K., Terner, J., Dawson, J. H. Caldariomyces fumago chloroperoxidase catalyzes the oxidative dehalogenation of chlorophenols by a mechanism involving two one-electron steps. J. Am. Chem. Soc. 129, 14838–14839 (2007).

50. Zambrano, G. et al. Oxidative dehalogenation of trichlorophenol catalyzed by a promiscuous artificial heme-enzyme. RSC Adv. 12, 12947–12956 (2022).

51. Zhu, H. et al. Tyrosinase-based proximity labeling in living cells and in vivo. J. Am. Chem. Soc., 146, 7515–7523 (2023).

52. Yoo, B. K. et al. Activation of p38 MAPK induced peroxynitrite generation in LPS plus IFN-γ-stimulated rat primary astrocytes via activation of iNOS and NADPH oxidase. Neurochem. Int. 52, 1188–1197 (2008).

53. Xia, Y., Zweier, J. L. Superoxide and peroxynitrite generation from inducible nitric oxide synthase in macrophages. Proc. Nat. Acad. Sci. 94, 6954–6958 (1997).

54. Garvey, E. P. et al. 1400W is a slow, tight binding, and highly selective inhibitor of inducible nitric-oxide synthase in vitro and in vivo. J. Biol. Chem. 272, 4959–4963 (1997).

55. Scott, G. S., Cuzzocrea, S., Genovese, T., Koprowski, H., Hooper, D. C. Uric acid protects against secondary damage after spinal cord injury. Proc. Nat. Acad. Sci. 102, 3483–3488 (2005).

56. Bylund, J., Brown, K. L., Movitz, C., Dahlgren, C., Karlsson, A. Intracellular generation of superoxide by the phagocyte NADPH oxidase: how, where, and what for? Free Radic. Biol. Med. 49, 1834–1845 (2010).

57. Amal, H. et al. S-nitrosylation of E3 ubiquitin-protein ligase RNF213 alters non-canonical Wnt/Ca+2 signaling in the P301S mouse model of tauopathy. Transl. Psychiatry 9, 44 (2019).

58. Thery, F. et al. Ring finger protein 213 assembles into a sensor for ISGylated proteins with antimicrobial activity. Nat. Commun. 12, 5772 (2021).

59. Okumura, F., Lenschow, D. J., Zhang, D.-E. Nitrosylation of ISG15 prevents the disulfide bond-mediated dimerization of ISG15 and contributes to effective ISGylation. J. Biol. Chem. 283, 24484–24488 (2008).

60. Takakura, K., Beckman, J. S., Ann MacMillan-Crow, L., Crow, J. P. Rapid and irreversible inactivation of protein tyrosine phosphatases PTP1B, CD45, and LAR by peroxynitrite. Arch. Biochem. Biophys. 369, 197–207 (1999).

61. Sevier, C. S., Qu, H., Heldman, N., Gross, E., Fass, D., Kaiser, C. A. Modulation of cellular disulfide-bond formation and the ER redox environment by feedback regulation of Ero1. Cell 129, 333–344 (2007).

